# Fine-scale proximity to offshore wind turbine foundations increases biomass of benthic fish species

**DOI:** 10.64898/2026.01.06.697906

**Authors:** Anthony W. J. Bicknell, Samuel Gierhart, Mario Lambrette, Matthew J. Witt

## Abstract

Offshore wind turbine fixed-bottom foundations provide artificial hard substrate through the water column that encourages marine flora and fauna to colonise and aggregate around the introduced structures, a well-documented phenomenon known as the ‘artificial reef effect’. The cumulative impact thousands of turbine foundations at multiple offshore sites have on local and regional marine species populations and communities is not fully understood. Knowledge of the extent and magnitude of the reefing effect at a fine scale (single turbines) is a prerequisite to making broader-scale (single or multiple wind farms) predictions of population level and ecosystem changes caused by presence of offshore wind farms. The influence of fine-scale distance (<250 m) to turbine jacket foundations on abundance, biomass and size of demersal fish was assessed at a northern latitude wind farm. Abundance and biomass of all demersal fish, flatfish *Pleuronectiformes spp.* and haddock *Melanogrammus aeglefinus* were found to have a significant negative relationship with increasing distance from foundations. Haddock were found to aggregate closer to the structures, yet all statistical models predicted a similar magnitude of increase for each group of between ∼1.5 and 1.6 times more individuals and biomass at 30 m from the foundations compared to 240 m. The results illustrate that fine-scale proximity to offshore wind fixed foundations has considerable effects on the presence of some demersal fish species. The cumulative or wider ecosystem consequences of these effects are not known, but the further evidence for localised reefing effects can be of strategic interest for optimizing future wind farm project design, included implementation of nature-inclusive measures that could help meet future marine net gain aspirations.

## 1 Introduction

Decarbonisation of global energy supply to mitigate the climate crisis has led to the development and growth of renewable energy sources on land and at sea (Østergaard et al., 2020). If internationally agreed emission targets are to be met (UNFCCC, 2015; Iacobuta et al., 2018) it will require a further rapid and extensive transition to low carbon energy sources with consideration for social (Carley and Konisky, 2020) and environmental impacts (Gayen et al., 2024).

Global offshore wind energy is a major part of the transition to renewable sources and is predicted to reach 55 GW capacity by 2034 (Williams et al., 2025), meaning a ∼6-7 fold increase in the next decade (8 GW = 2024 capacity), requiring the introduction of thousands more wind turbines and associated infrastructure to the marine environment (Díaz and Guedes Soares, 2020). The vast majority of the capacity (∼90%) is expected to be realised with wind turbines that are fixed to the seabed (‘fixed-bottom’) (Williams et al., 2025), which will be monopile, tripod, gravity based or jacket type foundations (Díaz and Guedes Soares, 2020). These structures introduce localised hard substrate habitat with vertical relief through the whole water column (sometimes complex, i.e. jackets) that are often colonised by sessile species (Degraer et al., 2020; Coolen et al., 2022) and attract mobile species looking for shelter and/or food (Langhamer, 2012; Buyse et al., 2022; Buyse et al., 2023a; Bicknell et al., 2025b), creating an ‘artificial reef effect’ (ARE) at each location (Langhamer, 2012).

The ARE is known to occur purposely or incidentally at subsea structures, such as designed artificial reefs (dos Santos et al., 2010), oil and gas platforms (Macreadie et al., 2011) or shipwrecks (Hickman et al., 2023), which can create highly productive fish habitats (Claisse et al., 2014; Smith et al., 2016; Birt et al., 2024). Studies focused on wind turbine foundations have found demersal fish species demonstrate residency behaviour (Reubens et al., 2013b; Buyse et al., 2023b) and have higher abundance and biomass in proximity to the structures (Reubens et al., 2013a; Reubens et al., 2014; Bicknell et al., 2025b), although it has not been determined conclusively whether the latter occurs through redistribution of existing fish or production of more (Reubens et al., 2013c). The design of wind farm foundation or seabed scour protection used can also influence the impact on fish occurrence (Werner et al., 2024; Bicknell et al., 2025b), making predictions of the effects across regional or global wind farm developments with varying foundation types and diverse fish communities a challenge. Studies where sampling has been close to turbine foundations (within 500 m) indicate that higher occurrence and biomass of demersal fish may diminish with increasing distance away from foundations (Methratta, 2021; Buyse et al., 2022) but this has not been formally assessed, although it has been shown for fish around introduced artificial reefs and oil and gas platforms (dos Santos et al., 2010; Egerton et al., 2021; Ibanez-Erquiaga et al., 2025).

Understanding whether, and to what extent (magnitude and spatial footprint), wind turbine foundations concentrate fish abundance is necessary to comprehend the effect large scale introduction of these structures will have on fish populations and the functioning of the wider ecosystem. For example, the concentration of fish biomass at subsea structures through either redistribution and/or production will change the prey energy landscape and behaviour of predators such as marine mammals (Russell et al., 2014; Fernandez-Betelu et al., 2022). For relatively small mammal species such as harbour porpoises *Phocoena phocoena* that have a high daily energy demands a change in availability and intake of food could have consequences for survival and reproduction (Booth, 2020). Moreover, details of the fine-scale effects offshore wind farm sites have on prey distribution could be used in models to estimate potential individual and population level effects on marine mammals that are known to forage in these areas (Booth et al., 2023). Such understanding and predictions of the impact a change in fish distribution could have on high priority species, such as marine mammals or seabirds, can help inform current and future consenting policy for wind farm developments and build evidence for No Net Loss (NNL) or Marine Net Gain (MNG) assessments (Hooper et al., 2021; Edwards-Jones et al., 2024).

Here we present a study to assess the effect of distance from wind turbine jacket foundations on the abundance, biomass and size of demersal fish at an offshore wind farm in Scotland, UK. Previous work at the wind farm revealed higher metrics in proximity (<50 m) to the jackets compared to references sites >500 m away (Bicknell et al., 2025b). By using a distance-based sampling design (Methratta, 2021) we test the hypothesis that the occurrence and size of demersal fish will decrease with increasing distance away from the foundation structures. The sampling was designed to focus on effects in a single year, rather than assessing interannual or seasonal variation in population, although it is known these could be significant (Bicknell et al., 2019).

## 2 Materials and Methods

### 2.1 Sampling Method and Equipment

Stereo Baited Remote Underwater Video (BRUV) was used to survey fish abundance, biomass and diversity in this study. The method is commonly used to survey fish (Harvey et al., 2021; McGeady et al., 2023), with advantages over many traditional survey methods, including; non-destructive, no or limited observer bias, data reanalysis possible and unrestricted depth (cost-dependent) (Bicknell et al., 2016). As with other survey methods it has bias, e.g. differentially attracting species, bait type, plume effects or restricted view, and these have been investigated elsewhere (e.g. Stobart et al., 2007). The BRUV systems used in this study were custom-made calibrated deepwater submersible rigs, each housing two cameras (Go-Pro Hero 9 Black) with LED lights mounted on the housing (see Bicknell et al., 2025b for details). The pole and bait box, which contained 100 g of frozen Atlantic mackerel *Scomber scombrus* (mackerel) bait, were attached to the housing and 15-20 kg of weight was fixed to the legs to temporarily secure the rigs to the seabed.

### 2.2 Study Location

The study took place in the Beatrice offshore wind farm in the Moray Firth, Scotland, North Sea (**Figure 1b**) between 15^th^-24^th^ July 2024 during daylight hours. The wind farm is located between ∼7.5-12 nautical miles offshore in water depth of between ∼35-60 metres. Beatrice operates 84 7MW turbines with jacket foundations that have 4 legs giving each a square with no scour protection. The jacket foundations were installed between August 2017 and July 2018. Commercial fishing is still permitted within the wind farm boundaries, with scallop dredging, trawling and creel potting known to take place (Dunkley and Solandt, 2022), sometimes close to the turbines (<50 m; pers comm). The seabed substratum across the wind farm comprised two biotopes: Medium to fine sands and muddy sands; or Cobbles and pebbles, gravels and coarse sands (Parry, 2019).

**Figure 1.**
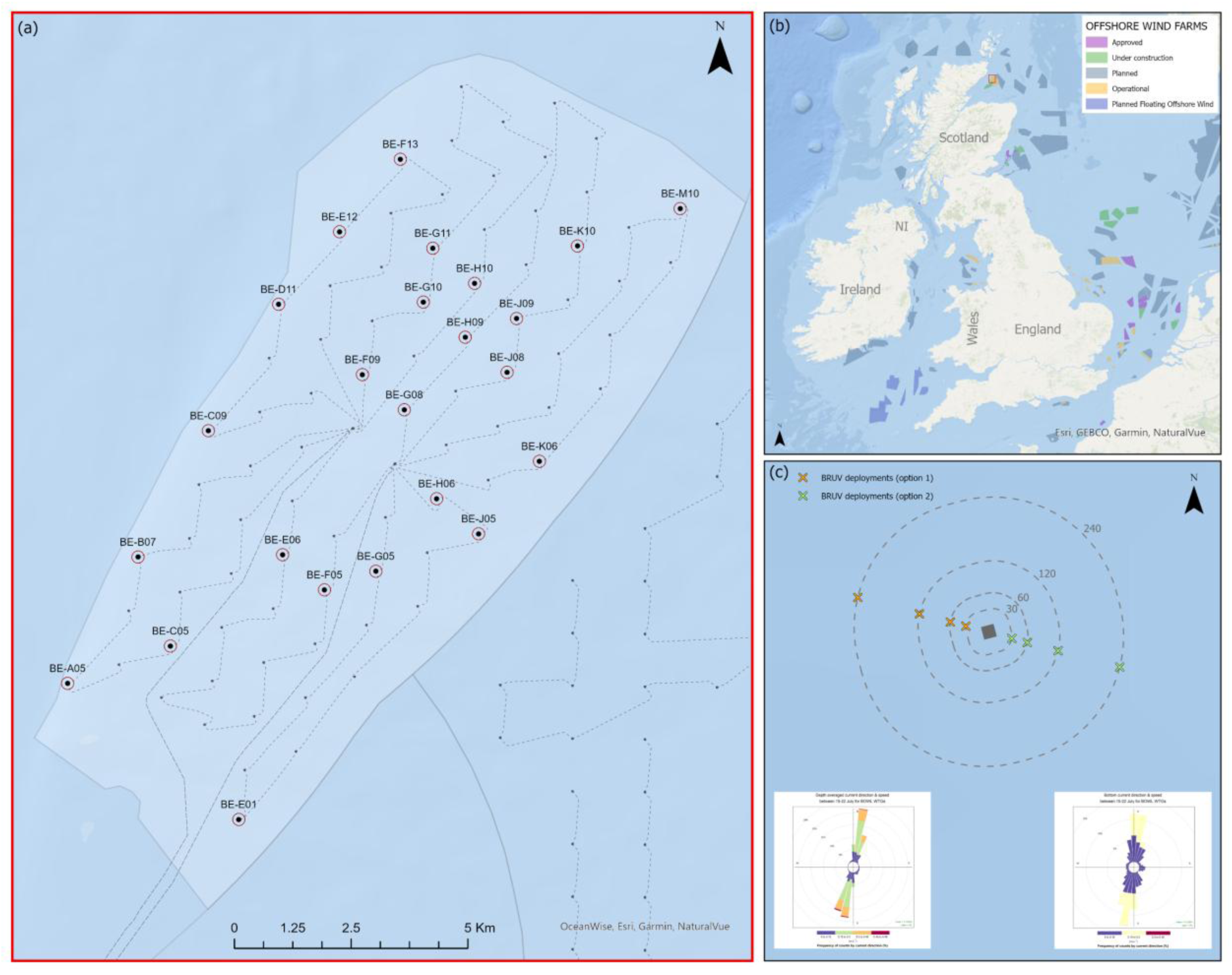
BRUV survey location and design. (a) The Beatrice offshore wind farm layout with turbine names and locations of the BRUV deployments. (b) The location of the Beatrice site and other operating or proposed wind farms around the British Isles and North Sea. (c) Illustrates the two options for deploying BRUVs at 4 distances from the wind turbine foundation, which are approximately perpendicular to the prevailing bottom and depth averaged current directions (see POLPRED data insets).

### 2.3 Survey and Design

BRUV systems were deployed at 30, 60, 120 and 240 m distances from 24 turbine jacket foundations within the Beatrice site (**Figure 1a**). The turbines for sampling were selected to provide two levels of density within the wind farm (high and low). The density for each turbine was calculated by counting the number of turbines that were located within a 2 km radius circle (**Supplementary Figure 1**). High density had 6 or more neighbouring turbines and low density had 3 or less. Four camera systems were simultaneously deployed to the seabed, one at each distance from the selected turbine foundation for 45 minutes before retrieval. BRUV were deployed in a line perpendicular to the prevailing current to avoid mooring entanglement risk and reducing bait plume overlap. Two deployment options (directions) were available at each turbine (WNW & ESE) and were chosen based on wind and weather conditions at the time to further reduce possible mooring entanglement (**Figure 1c**).

### 2.4 Image and Data Processing

The survey provided ∼72 hours of cumulative video data from 96 BRUV deployments, which were analysed using EventMeasure software (www.seagis.com.au). The analysis produced: number of demersal fish species, relative abundance (MaxN) and fish total body lengths (cm). MaxN is the maximum number of individuals, of each species, that occurred in a single video frame (MaxN frame). The use of MaxN to provide species relative abundance is widely used for BRUVs and is considered a conservative estimate of abundance, especially when species occur at a high density as no individual can be counted more than once (Willis and Babcock, 2000; Cappo et al., 2003).

Analysis of the video footage began once the BRUV landed on the seabed and the sediment settled and ended after 30 subsequent minutes (see Bicknell et al., 2019; Bicknell et al., 2025b). The relative abundance was determined by recording the first instance of a species and then subsequently only recording it again if the number of individuals for that species exceeded the previous MaxN. All individuals were identified to the lowest taxonomic level, apart from Common dab *Limanda limanda* and European flounder *Platichthys flesus,* which were grouped as *Pleuronectiformes spp.* (subsequently referred to as flatfish) since it was not possible to reliably distinguish between the two species on horizontal video footage. Evidence from previous studies and surveys in the Beatrice wind farm indicate common dab as the most likely and dominant species (Beatrice Offshore Windfarm Limited, 2021; Bicknell et al., 2025b).

MaxN estimates for haddock (*Melanogrammus aeglefinus*) and flatfish were created using a custom YOLO computer vision model (You Only Look Once; Ultralytics). Images (frames; n=2,709) were sampled from BRUV data collected at the survey location in 2022. These were annotated with bounding boxes in EventMeasure. Every fish in the frame was labelled with a close-fitting bounding box and its species name (*Pleuronectiformes spp.* for flatfish). Haddock and flatfish are known to be the dominant species in the local ecosystem (Bicknell et al., 2025b) and made up 80% of the bounding box annotations. The YOLO model was trained using only the haddock and flatfish annotations, resulting in a model that was able to detect both haddock and flatfish with high accuracy (**Supplementary Table 1**). The model was used to detect haddock and flatfish at 0.5 frames per second in all recorded survey footage (left- and right-camera) **(Video S1)**. Model detections were assigned a confidence score with optimal thresholds for including detections set to 0.4 for flatfish and 0.5 for haddock. The two camera angles showed variation within each deployment (**Supplementary Figure 2**), therefore, the MaxN for each deployment was taken as the highest abundance seen across the two viewpoints. During model development, validation was conducted on data during a 2022 survey at the site (**Supplementary Figure 3**). Once model results for the data presented here were obtained, performance was assessed on a random sample of 20 videos to ensure accuracy. These sample videos were manually analysed in EventMeasure and the MaxN estimates were compared to those produced by the YOLO model. Model accuracy was high and consistent with validation on previous data for both the left- and right-camera footage (**Supplementary Figure 4**).

Total body length (snout to tip of tail, cm) measurements were taken of individuals in the MaxN frame. Measurements of flatfish were taken manually using stereo images in EventMeasure, whereas length of the haddock were obtained using a semi-automated machine learning model in Automated Fish Identification (AFID), which is a custom plug-in for EventMeasure (Marrable et al., 2022). Haddock were labelled with a point in both the left and right videos which enables the AFID software to calculate body length. All AFID measurements were checked for accuracy and re-measured manually where necessary. A Residual Mean Square (RMS) value was calculated for each measurement, to indicate the error between corresponding measurement points in the stereo images (Marrable et al., 2022). Only measurements with RMS <20 were used in further analyses.

For individuals in a MaxN frame that could not be measured accurately (e.g. when animals were not sufficiently perpendicular to the BRUV rig and in a higher density scenario when fish bodies obscured each other), the mean length for that species MaxN frame was applied. In circumstances where there were no other individuals of that species in the video, the following hierarchy was followed until a mean measurement could be applied: within the same deployment; deployments at the same distance and density: deployments at the same distance.

To estimate individual fish weight the following standard equation was used:

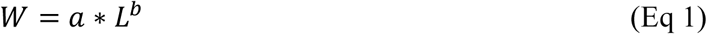

where W is the total weight in grams, L is the total length in cm (taken from the MaxN measurements), and **a** and **b** are species-specific conversion factors (Froese and Pauly, 2023).

The conversion factors were taken from the English North Sea ground fish survey (IBTS3E) for all fish species (Silva et al., 2013).

A biomass estimate for each deployment was then calculated using:

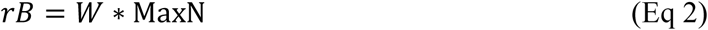

where *rB* is the total relative biomass in grams, W is the total weight in grams (Eq 1) and MaxN is the relative abundance.

The video footage were used to assign a habitat type code for each deployment: 1-rocky reef, 2-large coarse sediment, 3-medium mixed sediment, 4-fine sediment (as detailed in Bicknell et al., 2019). Habitat type was used in the subsequent analyses, alongside seabed current speed data (m/s) that was taken from POLPRED seabed CS20 models (https://noc-innovations.com/services/tide-prediction-software/offshore-software/).

### 2.5 Data Analysis

Relative abundance and biomass analyses were conducted for all demersal fish, flatfish and haddock *Melanogrammus aeglefinus*. Body length analyses were only conducted for flatfish and haddock *Melanogrammus aeglefinus* due to their high proportion of the total fish observed (cumulatively >94%; **Table 1**).

**Table 1.** Identified fish species on BRUVs; including number of deployments positively observed, number of individuals and the percentage of the overall total. * = High confidence of Common dab *Limanda limanda* but may contain European flounder *Platichthys flesus*.

| Common name | Scientific name | Family | Deployments (observed on) | Number of individuals | % of total species |
| --- | --- | --- | --- | --- | --- |
| *Flatfish | *Pleuronectiformes* spp. | *Pleuronectidae* | 96 | 1,687 | 64.2 |
| Haddock | *Melanogrammus aeglefinus* | *Gadidae* | 96 | 788 | 30.0 |
| Whiting | *Merlangius merlangus* | *Gadidae* | 38 | 64 | 2.4 |
| European plaice | *Pleuronectes platessa* | *Pleuronectidae* | 34 | 39 | 1.5 |
| Grey gurnard | *Eutrigla gurnardus* | *Triglidae* | 22 | 22 | 0.8 |
| Lesser spotted catshark | *Scyliorhinus canicula* | *Scyliorhinidae* | 12 | 12 | 0.5 |
| Flapper skate | *Dipturus intermedius* | *Rajidae* | 5 | 5 | 0.2 |
| Atlantic cod | *Gadus morhua* | *Gadidae* | 3 | 3 | 0.1 |
| Common dragonet | *Callionymus lyra* | *Callionymidae* | 1 | 2 | 0.08 |
| Goby sp. | *Pomatoschistus* sp. | *Gobiidae* | 1 | 1 | 0.04 |
| Spotted skate | *Raja montagui* | *Rajidae* | 1 | 1 | 0.04 |
| Skate sp. | *Raja* sp. | *Rajidae* | 1 | 1 | 0.04 |
| Poor cod | *Trisopterus minutus* | *Gadidae* | 1 | 1 | 0.04 |
| Total |  |  | 96 | 2832 | 100 |

To test for a relationship of proximity to turbine foundations generalized additive (GAMM) and generalized linear (GLMM) mixed models were fitted with distance as a smooth term, or fixed effect if the smooth term was not significant. Additionally, density of turbines (2-level factor), water depth, current direction (as viewed on camera: 4-level factor) and current speed were added as fixed effects and habitat type (3 levels) and turbine ID (24 levels) as random effects. The current direction term was assigned either North, South, East or West depending on the direction of the current relative to the camera (video footage). This was to incorporate any variability that might be caused by fish aggregating in the lee of the BRUV unit. Habitat type was not included as a fixed effect since it was not being directly tested and therefore not integrated in the survey design. When there was little variability resulting in a highly unbalanced random term, which can cause model instability, habitat was removed from models. Response data distributions were checked to determine model structure for each metric and comparison, resulting in a negative binomial family being used for abundance and gamma distribution used for biomass and length models. Model selection from all combinations of fixed effects in the full model was performed based on lowest AICc value (best fit) using the MuMin R package (Bartoń, 2023). Models with significant collinearity of fixed effect terms were removed from model selection. Model performance and diagnostics were performed using the residual simulation-based approach in the DHARMa R package (Hartig, 2022) and the performance package (Lüdecke et al., 2021). Analyses were conducted in R version 4.5.0 (R Core Team, 2025) using GAMLSS (Stasinopoulos and Rigby, 2007) and glmmTMB (Brooks et al., 2017) packages.

## 3 Results

### 3.1 Survey Sampling

A total of 96 BRUV deployments were completed (24 turbines x 4 distances; 100% of design: **Figure 1a**). There were 17 deployments that occurred on the NE side and 15 that occurred on the SW side of the turbine sites (**Figure 1c**). The depth range of deployments was 38-54 m, and current speeds were between 0.03-0.31 m/s. Three habitat types were observed on the BRUV footage: 82 (85.4%) fine sediment; 11 (11.5%) medium mixed sediment; (3.1%) large coarse sediment.

### 3.2 Image and Data Processing

A total of 2,475 length measurements were recorded for flatfish (n=1,687) and haddock (n=788). For flatfish, 1,120 (66.4%) lengths were directly measured, while 567 (38%) were derived from the mean measurements of the same deployment (n=566) or distance and turbine density (n=1). For haddock, 565 (71.7%) lengths were directly measured, and 223 (28.3%) were derived from the mean measurements of the same deployment (n=223).

### 3.3 Species Diversity and Taxonomic Composition

A total of 9 teleost and 3 elasmobranch species from 9 taxonomic families were observed by BRUVs across all deployments, representing 2,626 individual fish, the majority (>94%) were either flatfish or haddock (**Table 1**).

### 3.4 Proximity to turbine foundation (relative abundance and biomass)

The best fitting models to test distance to turbine for fish relative abundance and biomass contained linear terms, including depth and current direction (as viewed) (**Table 2**). Both metrics significantly decreased with increasing distance from turbine foundations (p <0.001; **Table 2**: **Figure 2**), with model estimates indicating ∼1.5 times more fish abundance and ∼1.47 times more biomass at 30 m compared to 240 m. Current direction to the south relative to the camera view was found to have significantly lower fish abundance and biomass (p<0.001; **Table 2**; **Supplementary Figure 5 & 6**), and there was a significantly negative relationship with increasing depth for biomass (p<0.05: **Table 2**; **Supplementary Figure 6**). No interaction terms were significant or retained in the best fitting models.

**Figure 2.**
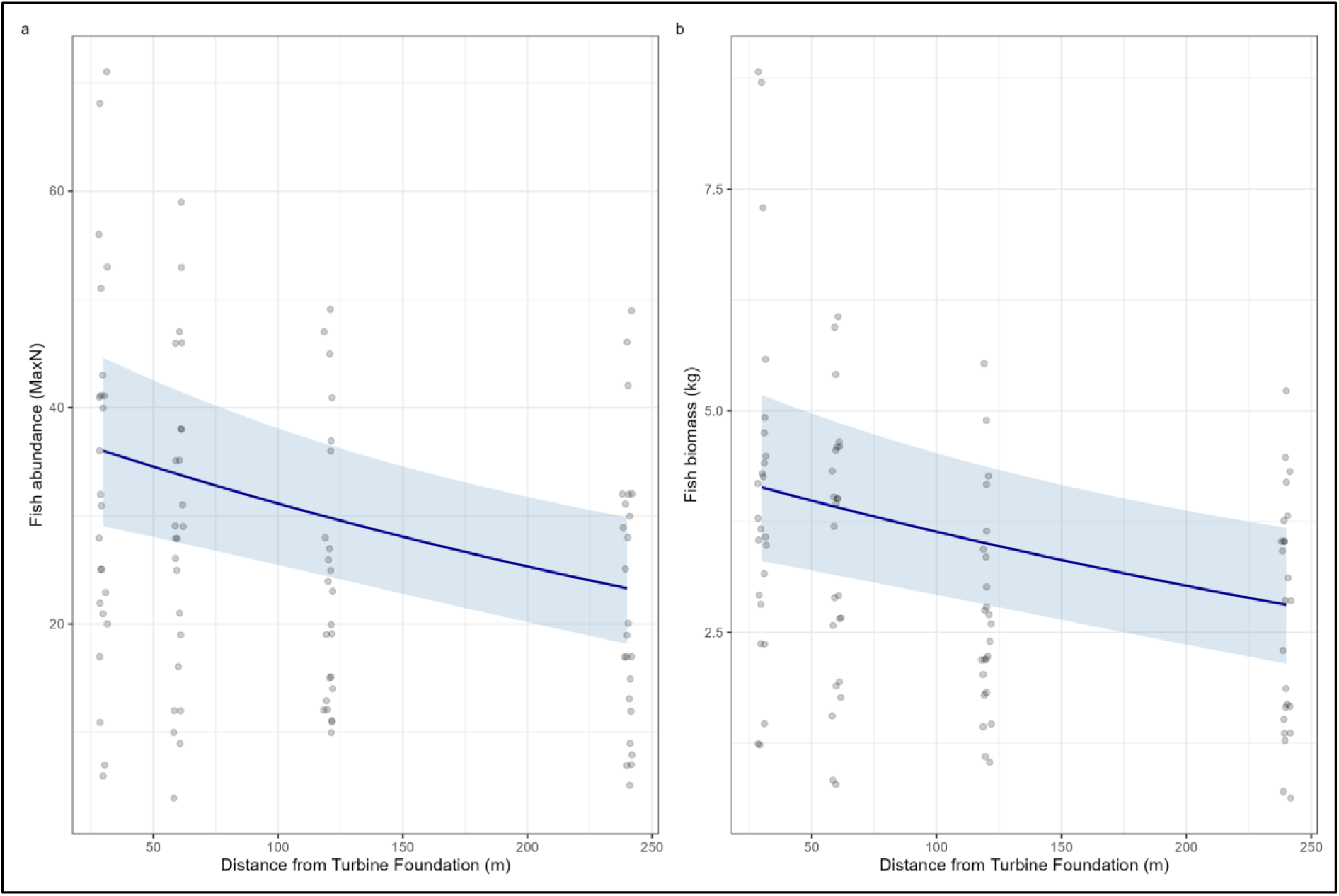
Fish abundance (a) and biomass (b) data with predicted GLMM estimate (± standard errors) for distance to turbine foundations. Grey circles = raw data points, blue line = model prediction, standard errors = shaded area.

**Table 2.** Best fitting generalized linear and additive mixed effect models for demersal fish, haddock and flatfish abundance and biomass. Terms with significant p-values below the 0.05 level are shown in bold.

|  | Fish abundance |  |  | Fish biomass |  |  | Flatfish abundance |  |  | Flatfish biomass |  |  | Haddock abundance |  |  | Haddock biomass |  |  |
| --- | --- | --- | --- | --- | --- | --- | --- | --- | --- | --- | --- | --- | --- | --- | --- | --- | --- | --- |
| **Fixed terms** | *Estimates* | *SE* | *p-values* | *Estimates* | *SE* | *p-values* | *Estimates* | *SE* | *p-values* | *Estimates* | *SE* | *p-values* | *Estimates* | *SE* | *p-values* | *Estimates* | *SE* | *p-values* |
| (Intercept) | 4.557 | 0.539 | **<0.001** | 9.468 | 0.441 | **<0.001** | 3.250 | 0.153 | **<0.001** | 7.077 | 0.197 | **<0.001** | 2.239 | 0.117 | **<0.001** | 8.641 | 0.512 | **<0.001** |
| Distance from Turbine | -0.002 | 0.001 | **<0.001** | -0.002 | 0.001 | **<0.001** | -0.002 | 0.001 | **0.001** | -0.002 | 0.001 | **0.017** |  |  |  |  |  |  |
| Depth | -0.020 | 0.012 | 0.089 | -0.023 | 0.009 | **0.012** |  |  |  |  |  |  |  |  |  | -0.019 | 0.011 | 0.088 |
| Current (North) | 0.092 | 0.118 | 0.431 | 0.021 | 0.135 | 0.876 | 0.076 | 0.145 | 0.601 | 0.074 | 0.205 | 0.718 | -0.100 | 0.134 | 0.457 |  |  |  |
| Current (South) | -0.878 | 0.166 | **<0.001** | -0.668 | 0.148 | **<0.001** | -1.141 | 0.219 | **<0.001** | -1.141 | 0.231 | **<0.001** | -0.624 | 0.165 | **<0.001** |  |  |  |
| Current (West) | -0.217 | 0.126 | 0.084 | -0.111 | 0.136 | 0.416 | -0.364 | 0.157 | **0.021** | -0.429 | 0.218 | **0.049** | -0.108 | 0.140 | 0.441 |  |  |  |
| Turbine density (High/Low) |  |  |  |  |  |  |  |  |  |  |  |  |  |  |  | -0.218 | 0.108 | **0.046** |
| **Smooth terms** |  |  |  |  |  |  |  |  |  |  |  |  | *EDF* | *Chi. sq* | *p-values* | *EDF* | *F* | *p-values* |
| s(Distance from Turbine) |  |  |  |  |  |  |  |  |  |  |  |  | 1.793 | 15.08 | **<0.001** | 1.557 | 4.184 | **0.001** |
| s(WTG)<sub>random</sub> |  |  |  |  |  |  |  |  |  |  |  |  | 10.965 | 21.08 | **0.005** | 2.026 | 0.117 | 0.216 |
| **Random Effects** |  |  |  |  |  |  |  |  |  |  |  |  |  |  |  |  |  |  |
| $\sigma^2$ | 0.309 | | | 0.158 | | | 0.340 | | | 0.309 | | | | | | | | |
| $\tau_{00}$ | 0.031<sub>WTG</sub> | | | 0.001<sub>WTG</sub> | | | 0.134<sub>WTG</sub> | | | 0.129<sub>WTG</sub> | | | | | | | | |
| ICC | 0.091 |  |  | 0.005 |  |  | 0.283 |  |  | 0.295 |  |  |  |  |  |  |  |  |
| N | 24<sub>WTG</sub> |  |  | 24<sub>WTG</sub> |  |  | 24<sub>WTG</sub> |  |  | 24<sub>WTG</sub> |  |  |  |  |  |  |  |  |
| Observations | 96 |  |  | 96 |  |  | 96 |  |  | 96 |  |  | 96 |  |  | 96 |  |  |
| Marginal R<sup>2</sup> / Conditional R<sup>2</sup> | 0.313 / 0.376 |  |  | 0.383 / 0.386 |  |  | 0.327 / 0.517 |  |  | 0.338 / 0.534 |  |  | 0.376 |  |  | 0.180 |  |  |
| Model type | GLMM |  |  | GLMM |  |  | GLMM |  |  | GLMM |  |  | GAMM |  |  | GAMM |  |  |
| Family | Negative binomial |  |  | Gamma |  |  | Negative binomial |  |  | Gamma |  |  | Negative binomial |  |  | Gamma |  |  |

The best fitting models to test distance to turbine for flatfish relative abundance and biomass contained linear terms, including current direction (as viewed) (**Table 2**). Both flatfish metrics significantly decreased with increasing distance from turbine foundations (p <0.05; **Table 2**: **Figure 3**), with model estimates indicating ∼1.6 times more flatfish abundance and ∼1.6 times more biomass at 30 m compared to 240 m. Current direction to the south relative to the camera view was found to have significantly lower flatfish abundance and biomass (p<0.001; **Table 2**; **Supplementary Figure 7 & 8**). No interaction terms were significant or retained in the best fitting models.

**Figure 3.**
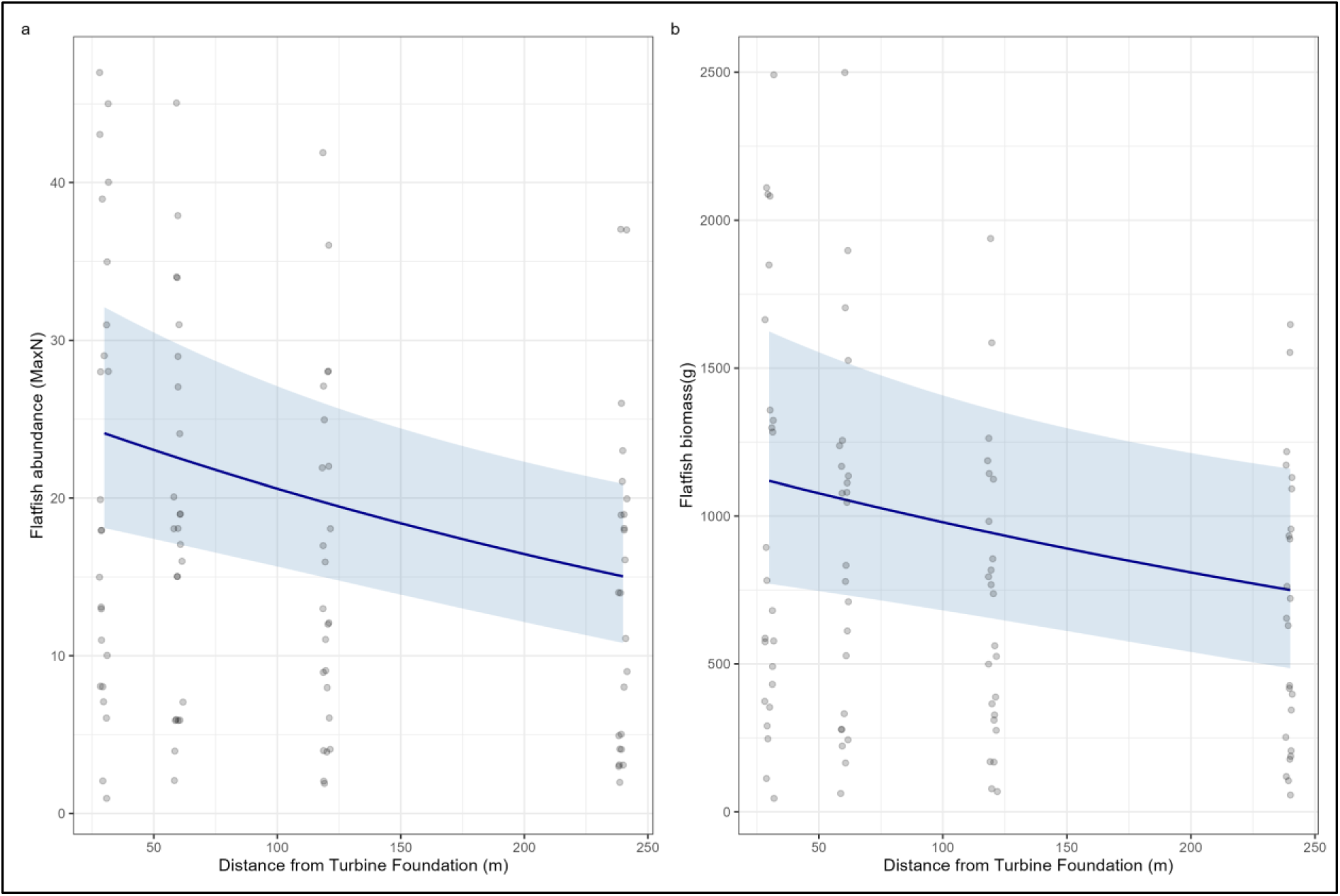
Flatfish abundance (a) and biomass (b) data with predicted GLMM estimate (± standard errors) for distance to turbine foundations. Grey circles = raw data points, blue line = model prediction, standard errors = shaded area.

The best fitting model to test distance to turbine for haddock relative abundance contained a smooth term for distance, and linear term for current direction (as viewed) (**Table 2**). The best model for haddock biomass contained a smooth term for distance, and linear terms for depth and turbine density (**Table 2**). Both metrics significantly decreased with increasing distance from turbine foundations (p <0.001; **Table 2**: **Figure 4**), with model estimates indicating ∼1.5 times more haddock abundance and biomass at 30 m compared to 240 m. Current direction to the south relative to the camera view was found to have significantly lower haddock abundance (p<0.001; **Table 2**; **Supplementary Figure 9**). Haddock biomass was found to be significantly higher (∼1.2 times) at turbines in high density areas of the wind farm (p<0.05: **Table 2**; **Supplementary Figure 10**). No interaction terms were significant or retained in the best fitting models.

**Figure 4.**
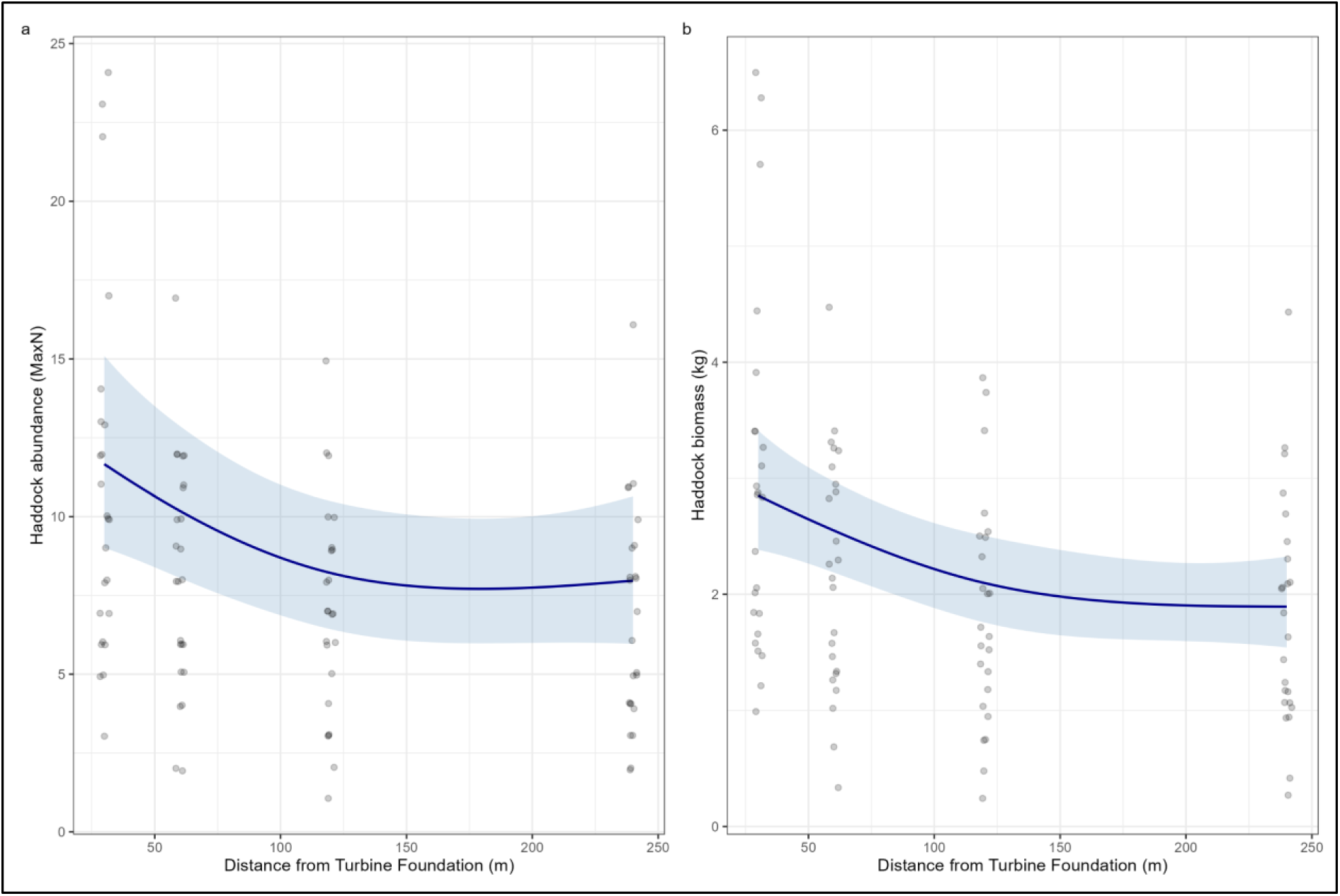
Haddock abundance (a) and biomass (b) data with predicted GAMM estimate (± standard errors) for distance to turbine foundations. Grey circles = raw data points, blue line = model prediction, standard errors = shaded area.

### 3.5 Proximity to turbine foundation (flatfish and haddock size)

The best fitting model for flatfish length was the null model with no effect of distance from turbines (**Supplementary Figure 11b**). For haddock, distance from turbines was also not significant (**Supplementary Figure 11a**) and only depth was retained in the best fitting linear model (**Table S2**). Body length significantly decreased with increased depth (p<0.05; **Supplementary Table 2**; **Supplementary Figure 12**).

## 4 Discussion

Abundance and biomass of demersal fish were found to decrease with increasing distance from jacket foundations in summer 2024 at the Beatrice offshore wind farm in the Moray Firth, Scotland. The effect was a linear decrease for all demersal fish or flatfish but showed a decreasing gradient for haddock with a steeper slope closer to the structures (<60 m). The shape and rate of decline with increasing distance from turbine foundations varied but the magnitude was similar for all demersal fish, flatfish and haddock with between ∼1.5 and 1.6 times more individuals and biomass estimated at 30 m from the foundations compared to 240 m. There was no significant change in the body length of haddock or flatfish with fine-scale (<240m) distance from the turbine foundations.

### 4.1 Species and communities

The demersal teleost fish species observed (n = 9) around the Beatrice turbine foundations were a subset of the species found in a previous survey using the same technique inside and outside the Moray Firth wind farms in 2022 (n=13: Bicknell et al., 2025b), and using trawling for post-construction surveys in 2021 (n=12-13: Beatrice Offshore Windfarm Limited, 2021). There was a similar heavy skew of over 90% of the abundance being flatfish (common dab) and haddock as with the 2022 survey (Bicknell et al., 2025b). However, there was a higher proportion of haddock found in the 2024 survey (∼30%) when compared to the more spatially extensive survey in 2022 (∼20%), which included reference sites further from the turbines (500 m) and outside wind farm boundaries (>2 km from turbines). The proportional increase aligns with more focused sampling close to turbine foundations in 2024 and the knowledge from both studies that haddock aggregate in greater numbers close to these jacket structures in the Moray Firth.

### 4.2 Proximity to turbine foundation (abundance, biomass and size)

The observed and modelled decline of demersal fish abundance and biomass with increasing distance from turbine foundations supported the *a priori* hypothesis and provides further evidence for the localised nature (<100 m) of the ARE on demersal fish distribution around foundations (Bergström et al., 2013; Methratta, 2021; Buyse et al., 2022; Werner et al., 2024; Bicknell et al., 2025b). Increased fish abundance with proximity to subsea structures has also been observed at introduced artificial reefs (dos Santos et al., 2010; Rosemond et al., 2018) and offshore oil and gas platforms (Reynolds et al., 2018; Egerton et al., 2021) with recent studies around platforms in the Danish North Sea finding fine-scale (<100 m) increase in abundance for demersal gadoid and flatfish species similar to the current study (Ibanez-Erquiaga et al., 2025). Studies have consistently shown higher abundance close to turbine foundations (jacket or monopiles) of species that are known to associate with hard bottom, complex environments compared to the surrounding mobile seabed (e.g. Atlantic cod Gadus morhua, pouting Trisopterus luscus, Shorthorn sculpin Myoxocephalus scorpius; Bergström et al., 2013; Reubens et al., 2013a; van Hal et al., 2017). Based on the soft or mobile sediment preference of adult haddock, common dab and other small flatfish species (Brodziak, 2005; Able and Joel Fodrie, 2014) these species may not be expected to aggregate around these structures even when present in the wind farm. There is growing evidence, however, to suggest this behaviour is not uncommon in these species across foundation types and locations (Buyse et al., 2022; Bicknell et al., 2025b; Ibanez-Erquiaga et al., 2025), and studies at scour protected monopiles in Belgium linked this behaviour to food availability (Buyse et al., 2023a; Buyse et al., 2023b) and/or shelter for flatfish (Buyse et al., 2022). With fixed foundation offshore wind farms typically built on soft sediment shallow shelf sea environments, such as the North Sea in Europe (Gierhart et al., 2024) and the Yellow or China Seas in Asia (Zhang and Wang, 2022), demersal fish with close association with these conditions (<100 m depth and soft bottom) are disproportionately impacted by loss of suitable habitat when its replaced with turbine foundations, scour protection and cabling infrastructure. The direct loss of habitat for flatfish species appears to be balanced with indirect effects of enhanced food availability through the creation of 3-dimensional, hard substrate habitat that supports increased abundance and diversity of prey organisms (Reubens et al., 2014; Coolen et al., 2022) and potential refuge from predation (Buyse et al., 2022). The same may be concluded for haddock based on the close aggregation (<30 m) to turbine foundations found in the Beatrice wind farm, but further research targeting prey consumption (e.g., stomach content, stable isotope analyses) and detailed movement behaviour at this and other wind farm locations would be needed to provide relevant evidence.

The lack of a significant change in body length with increasing distance from turbine foundations for either haddock or flatfish indicates that at the spatial scale assessed in this study there was little variation in the year classes during summer 2024. Significant size differences were found at larger spatial scales (∼500 m & 2 km) within and around Beatrice wind farm in 2022, which equated to a larger proportion of 3 and 4 year old haddock and flatfish at turbine foundations and inside the wind farm site (Bicknell et al., 2025b). When size and estimated age from the samples collected 30 m from turbines were compared between the two survey years (age estimation described in Bicknell et al., 2025b) they differ (larger haddock and smaller flatfish in 2024: **Supplementary Figures 13-15**), but these data are not useful to determine trends and are only illustrative of annual variation inherent in fish populations, which was not examined in these studies. To relate these data to commercial fisheries in the UK North Sea, the size distribution across both 2022 and 2024 for haddock indicate approximately half (52%) observed within 30 m of the foundations where above minimum landing size for the UK (30 cm: Minimum Conservation Reference Sizes (MCRS) in UK waters - GOV.UK). Haddock is a major part of the “Big 5” commercially exploited species in Europe (referring to cod, haddock, salmon, tuna, and prawns/shrimp) so of value to the UK seafood industry (Heard et al., 2025). Although dab (or flounder) does not have catch quotas in Europe there is a minimum landing requirement of 15 cm in the UK, which >75% of the measured flatfish where above. Knowledge of how individual or multiple OWFs alter distribution, biomass or age structure could be valuable for commercially landed fish with catch limits that are derived from scientific monitoring surveys and fisheries catch data at regional scales (ICES, 2025).

Scientific research surveys and trawl fishing are permitted inside UK wind farms but these activities are perceived to be challenging due to safety concerns related to ship and gear proximity to turbine foundations and other subsea infrastructure (Gray et al., 2016), resulting in less trawling within certain OWFs (Dunkley and Solandt, 2022). These limitations or exclusion of survey and fishing activity within OWFs have led to concerns as to how these sites will affect long term monitoring programs that are essential for fish stock assessment and setting catch limits for management (Hogan et al., 2023; Gill et al., 2025). Moreover, if surveys or trawl fishing take place in OWFs they are likely to occur at a distance from the structures, and this could mean important spatial structure is missing. Additional influence on the accuracy of the survey data used in stock modelling may arise if the fine-scale biomass aggregation (<100 m) and age structure changes (<500 m) found for haddock at Beatrice and Moray East wind farm are general effects across demersal species, wind turbines foundations and sites. Integrating data from diverse sources and application of spatially explicit modelling techniques could help account for these fine scale effects of OWFs for commercially important species such as haddock (Cao et al., 2020; Silva et al., 2025), but would require incorporation into existing national and international long-term monitoring programs and stock modelling policies (e.g. ICES, 2025).

While turbine foundations were found to increase haddock and flatfish abundance and biomass at a fine scale, it may not constitute a net positive for the populations if the fish are drawn from elsewhere and not from enhanced production of new biomass (Gill et al., 2020; Czaja Jr et al., 2025). Direct evidence for enhanced production (or not) is required to assess the ultimate ecosystem consequences wind farms have when altering distribution and/or demographics of demersal fish populations (Gill et al., 2025). Indirect evidence indicate potential higher production for demersal species through increased feeding opportunities (Reubens et al., 2013c; Buyse et al., 2023a) and reproduction (Gimpel et al., 2023), but direct evidence through quantitative estimates needs to be an aim of future studies (Czaja Jr et al., 2025). Evidence of enhanced fish production at offshore oil and gas infrastructure (Claisse et al., 2014; Birt et al., 2024) and artificial reefs (Smith et al., 2016) provide a good indication that it could also occur at wind turbines over time given the similar reefing effects. These potential responses lead to the challenging question as to the cumulative impact of thousands of turbines across multiple wind farms on fish populations and the wider communities.

### 4.3 Wider ecosystem impacts

The widespread installation of offshore wind turbine structures will inevitably alter habitat and species population connectivity across large areas progressing from a localised to a seascape-level phenomenon (Isaksson et al., 2023). Through provision of new habitat and change in food availability, an inter-connected network of artificial reefs has the potential to alter the dynamics of fish populations with subsequent ecosystem effects (Isaksson et al., 2023). Whether attraction and aggregation of fish creates ecological traps and ‘sink’ populations (Kristan, 2003; Reubens et al., 2013c), or leads to highly productive ‘source’ populations (e.g. Claisse et al., 2014; Birt et al., 2024), the effects are likely to be species specific. A significant challenge is to robustly predict how these species level effects will scale up to meta-populations and wider ecosystem functioning given the amount of synchronous spatio-temporal empirical data required to parameterize large meta-ecology models effectively (e.g. life-history, life stage dispersal, mobility, genetic populations: Warmuth et al., 2025).

Relatively small scale ecosystem models have predicted that changes in the distribution and/or production (biomass) of fish caused by turbine foundations can change local food-webs, with positive effects for upper trophic levels such as marine mammals and seabirds (Raoux et al., 2017). Altering the location, amount, availability and/or nutritional quality of food (energy) influences behaviour and daily energy intake of individual animals (predators), which can have consequences for their fitness (Nabe-Nielsen et al., 2018; Booth, 2020; Booth et al., 2023). Individual or agent-based models have been developed to simulate how these changes in vital rates may lead to population-level impacts (Mortensen et al., 2021). For marine mammals and seabirds, models such as iPCoD (King et al., 2015), DEPONS (Nabe-Nielsen et al., 2018) and SeabORD (Searle et al., 2018) have been developed and/or being applied to assess long-term population-level effects of offshore wind farm disturbances, including sound exposure during construction, displacement and change in prey densities. These models are being developed further to incorporate additional spatio-temporal empirical prey (fish) data to provide more accurate estimations for use in wind farm Environmental Impact Assessments (EIA) and consenting requirements (see the PrePARED Project [PrePARED – An offshore renewables science project](https://owecprepared.org/)). The acquisition and inclusion of empirical data on changes in fish distribution, biomass and behaviour (e.g. Bicknell et al., 2025a) caused by OWFs is necessary to reduce inherent uncertainties in ecosystem and individual based models.

### 4.4 Marine net gain and nature-inclusive design

The prevailing regulatory paradigm is on minimizing harm and achieving no net loss (NNL) of biodiversity following development projects, such as offshore wind (zu Ermgassen et al., 2019). However, the emerging and more ambitious concept of marine net gain (MNG) seeks to leave the marine environment in a measurably better state than before development (Hooper et al., 2021). In simple terms, MNG will require an amount of gain to be proportional to the magnitude of residual loss of relevant environmental features associated with each project (DEFRA, 2022). While MNG frameworks and metrics are still being established, a sophisticated understanding of how biodiversity and ecosystems are affected by wind farm developments is key for practical and effective application (Hooper et al., 2021; Edwards-Jones et al., 2024). For example, the UK government is considering plans to extend MNG assessments to species (as well as habitats) and beyond the footprint of the development to help consider effects at the ecosystem level, but possibly exclude artificial reef and other relevant potentially positive incidental impacts due to the uncertainty of the benefits (DEFRA, 2022; 2023).

In the case of Beatrice OWF, it has been found that the reefing effect at operating foundations can redistribute the locally dominant demersal fish species at a fine-scale, which could possibly lead to enhanced population production and likely changes in predator foraging behaviour (e.g. marine mammals; Fernandez-Betelu et al., 2022). Although there is uncertainty in the consequences for local populations, predators and the wider ecosystem (positive, negative or neutral) it would seem remiss for reefing effects such as these not to be considered and accounted in policy when they are established and evidenced impacts. Proposals and use of nature-inclusive foundation designs or materials that maximise ecological benefits illustrate how confidence in AREs are influencing wind farm mitigation measures (Hermans et al., 2020; Kingma et al., 2024). Nature-inclusive design has the potential to benefit target habitats or species, such as ‘cod hotels’ for Atlantic cod (Hermans et al., 2020), but if incidental reef effects are not incorporated into MNG assessments the benefits would need to be considered relative to baseline levels (incidental effects) to evaluate the additional gain and incentivise use of a design.

## 5 Conclusion

The effect of offshore wind foundations on demersal fish can create significant fine-scale local change by aggregating fish and potentially produce new biomass, but whose cumulative impacts and effect on the wider ecosystem are complex and not yet fully understood. For wind farm developers, knowledge of these fine-scale, site and foundation specific processes can be of strategic interest for optimizing project design including nature-inclusive measures. By strategically harnessing ARE and proactively managing its cumulative consequences, the offshore wind industry has the potential to become not just a source of clean energy, but a significant driver of marine restoration and ecosystem gain.

## Data availability statement

Data available via designated repository: (*tbc*)

## Ethics statement

All study activities were conducted in compliance with the University of Exeter’s ethics and Health and safety policies.

## Author contributions

A.W.J. Bicknell: Conceptualisation, Funding acquisition, Investigation, Data curation, Formal analysis, Visualization, Writing - Original Draft. S. Gierhart: Investigation, Data curation, Writing – review & editing. M. Lambrette: Investigation, Data curation, Formal analysis, Writing – review & editing. M.J. Witt: Conceptualisation, Funding acquisition, Writing – review & editing.

## Acknowledgements

Thanks to the funders, PrePARED project partners, the skipper and crew of the vessel Waterfall (Moray First Marine Ltd).

