## Supplementary Material for "Fine-scale proximity to offshore wind turbine foundations increases biomass of benthic fish species"

### 1 Supplementary Figures and Tables

#### 1.1 Supplementary Figures

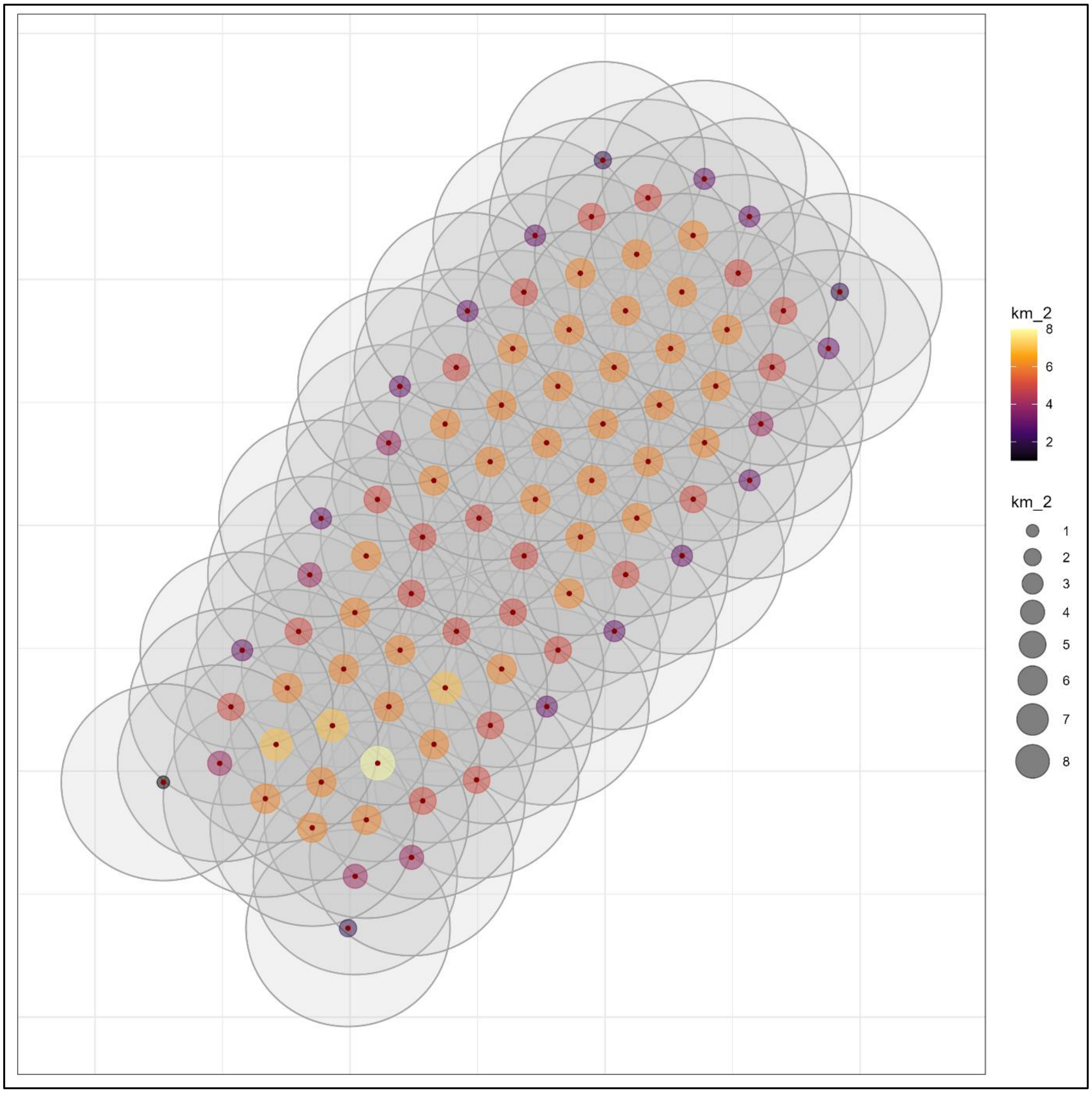


**Supplementary Figure 1.** Spatial plot of turbine density at a 2 km radius scale. Dots = turbine foundations, grey shaded circles = 2 km radius around each turbine, filled coloured circles = density (number of turbines within 2 km radius) shown by size and colour as per legends.

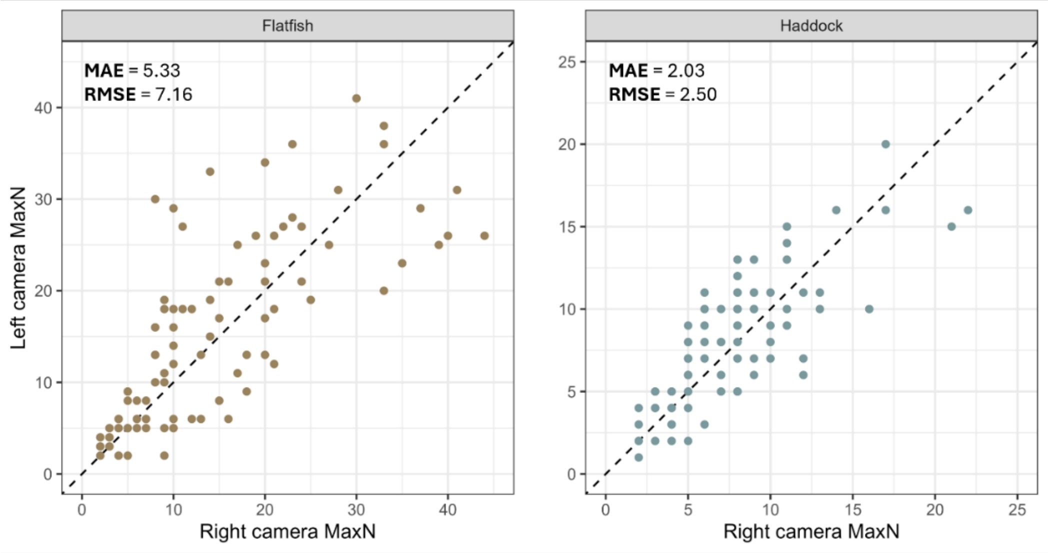


**Supplementary Figure 2.** MaxN estimates from the left and right camera of the stereo-BRUV systems for haddock and flatfish. The root mean square error (RMSE) and mean absolute error (MAE) for right camera relative to the left are shown.

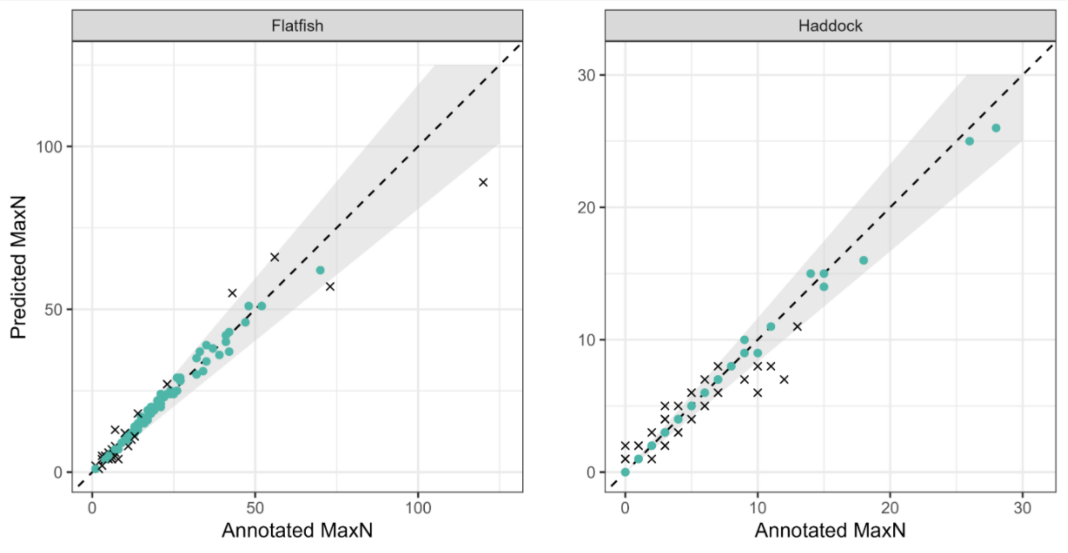


**Supplementary Figure 3.** Comparison between YOLO model-derived and human-annotated MaxN on survey footage collected at the study location in 2022. This data was used for model training, development and validation. The shaded area shows the bounds around perfect agreement which we deemed 'acceptable' for model validation purposes.

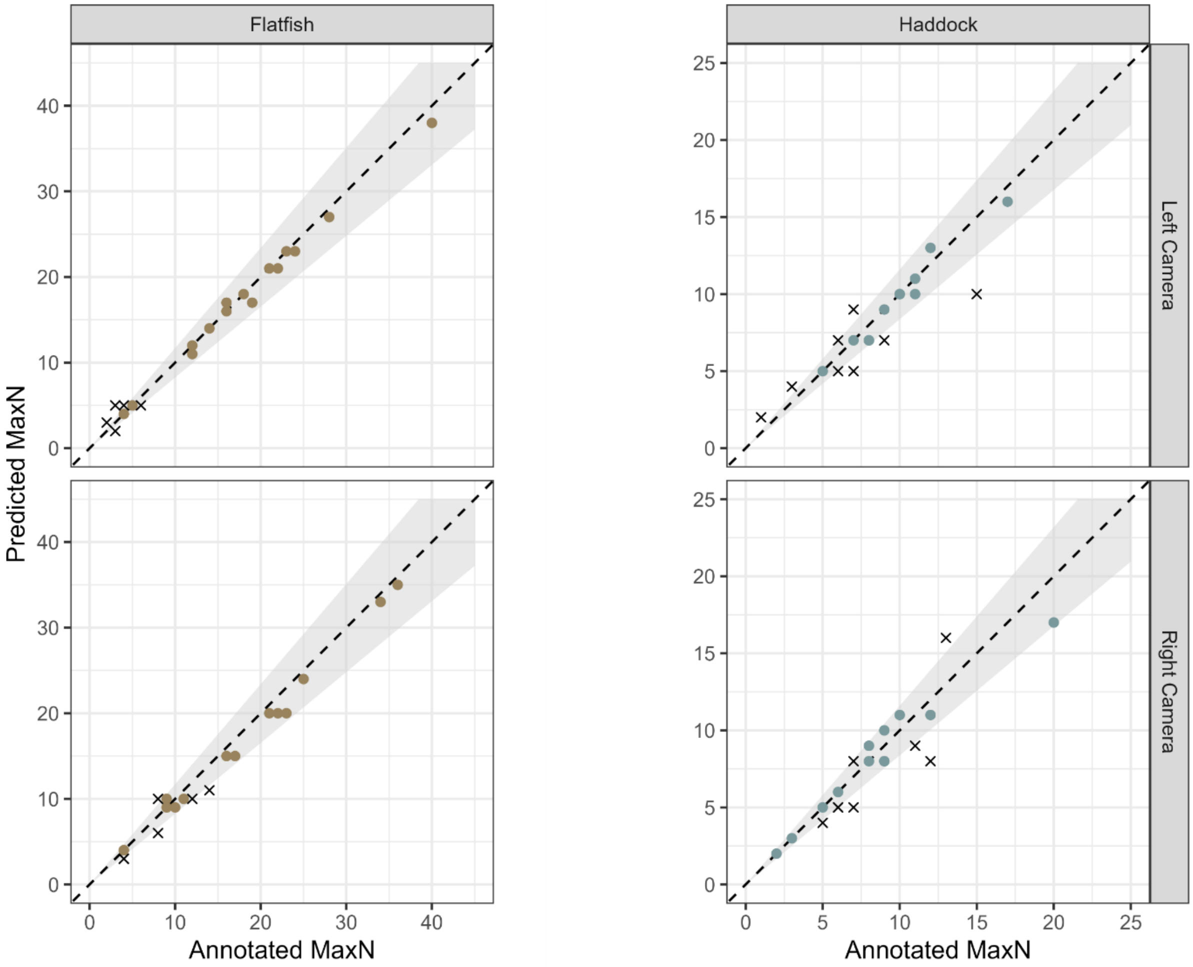


**Supplementary Figure 4.** Comparison between YOLO model-derived and human-annotated MaxN on 20 randomly selected deployments from this study. The shaded area represents the same bounds for 'acceptable' error as those shown in **Supplementary Figure 3** Model performance on these deployments was consistent with performance on the 2022 validation data.

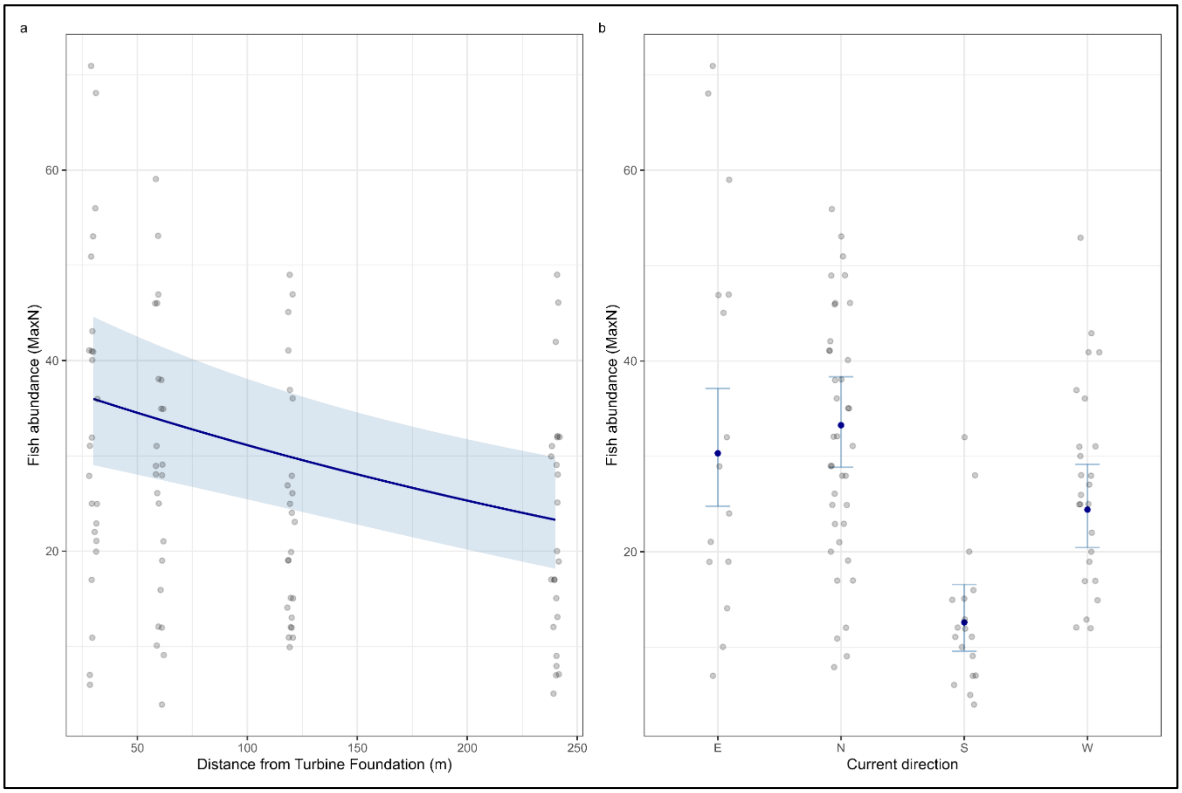


**Supplementary Figure 5.** Plots of raw data and estimated values with standard errors (light blue shaded ribbon or error bars) for significant terms from the demersal fish abundance GLMM.

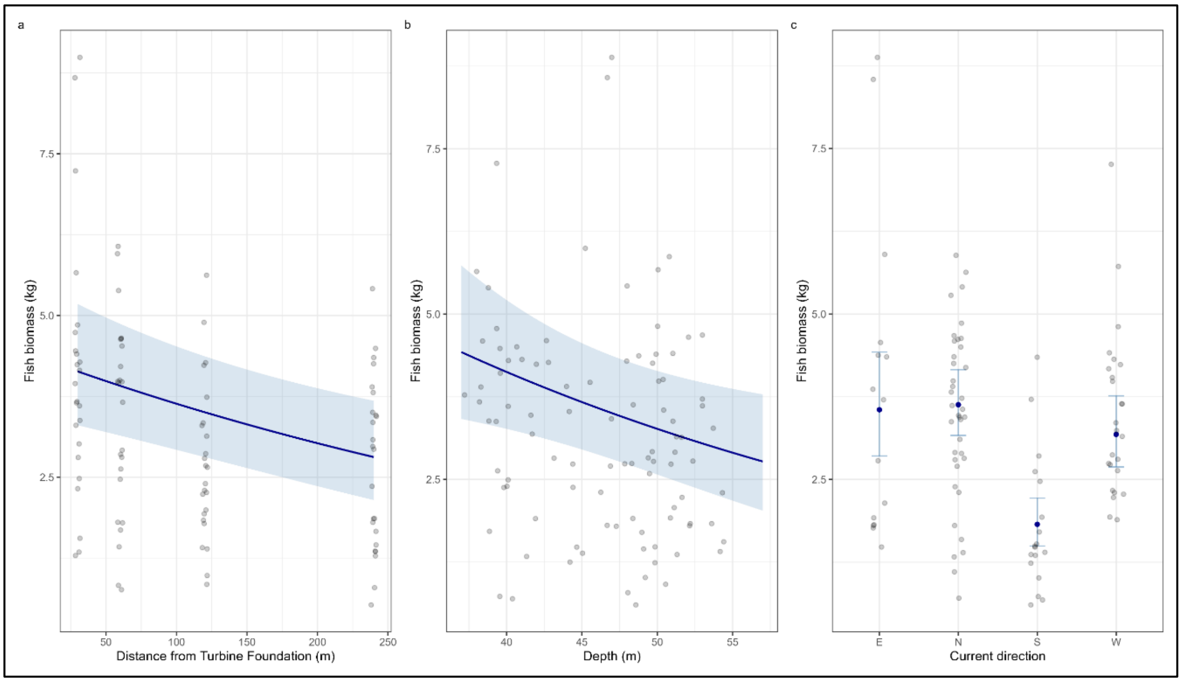


**Supplementary Figure 6.** Plots of raw data and estimated values with standard errors (light blue shaded ribbon or error bars) from significant terms from the demersal fish biomass GLMM.

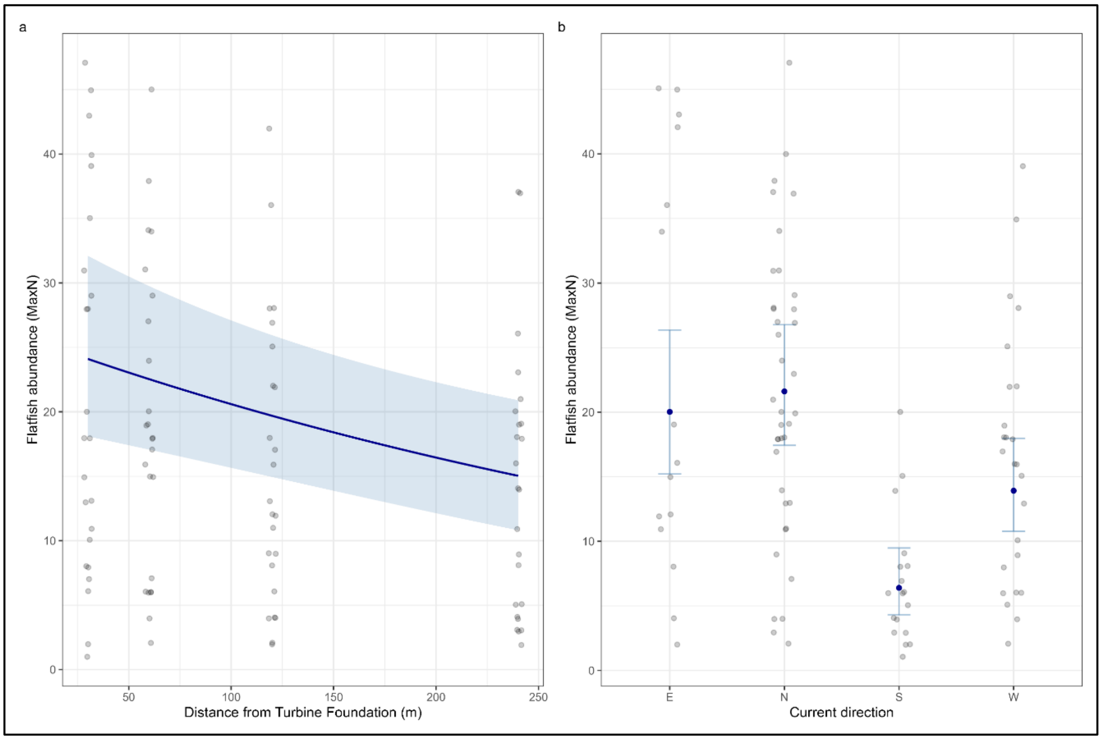


**Supplementary Figure 7.** Plots of raw data and estimated values with standard errors (light blue shaded ribbon or error bars) for the flatfish abundance GLMM.

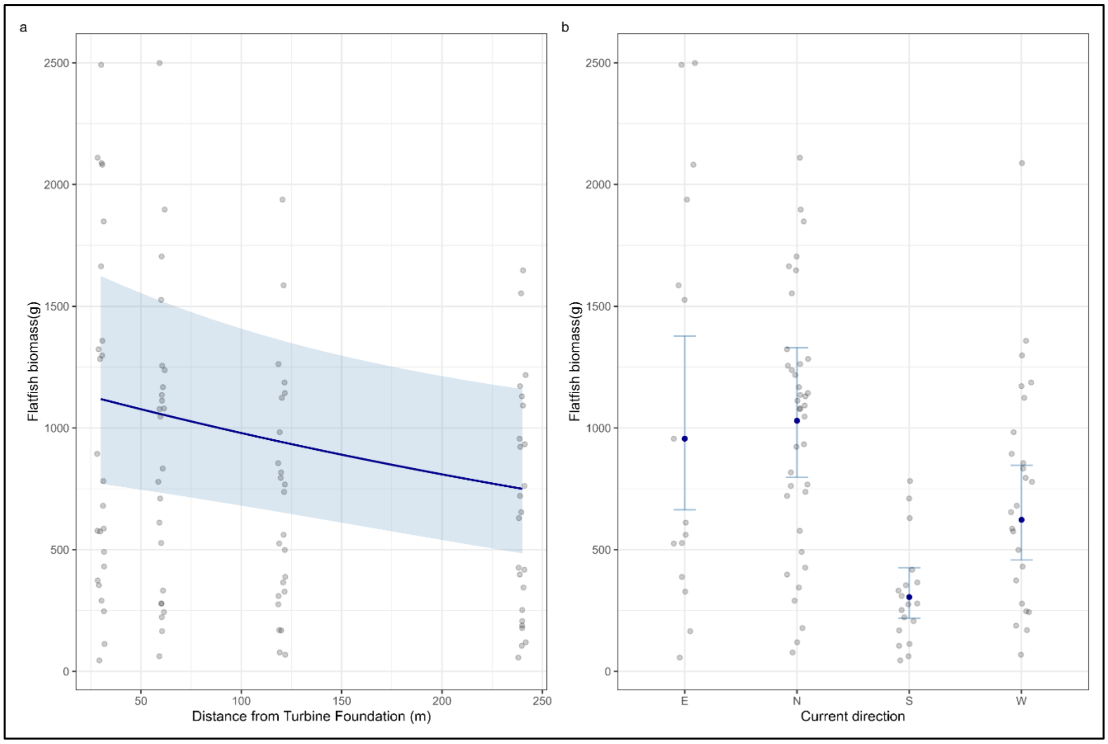


**Supplementary Figure 8.** Plots of raw data and estimated values with standard errors (light blue shaded ribbon or error bars) for significant terms from the flatfish biomass GLMM.

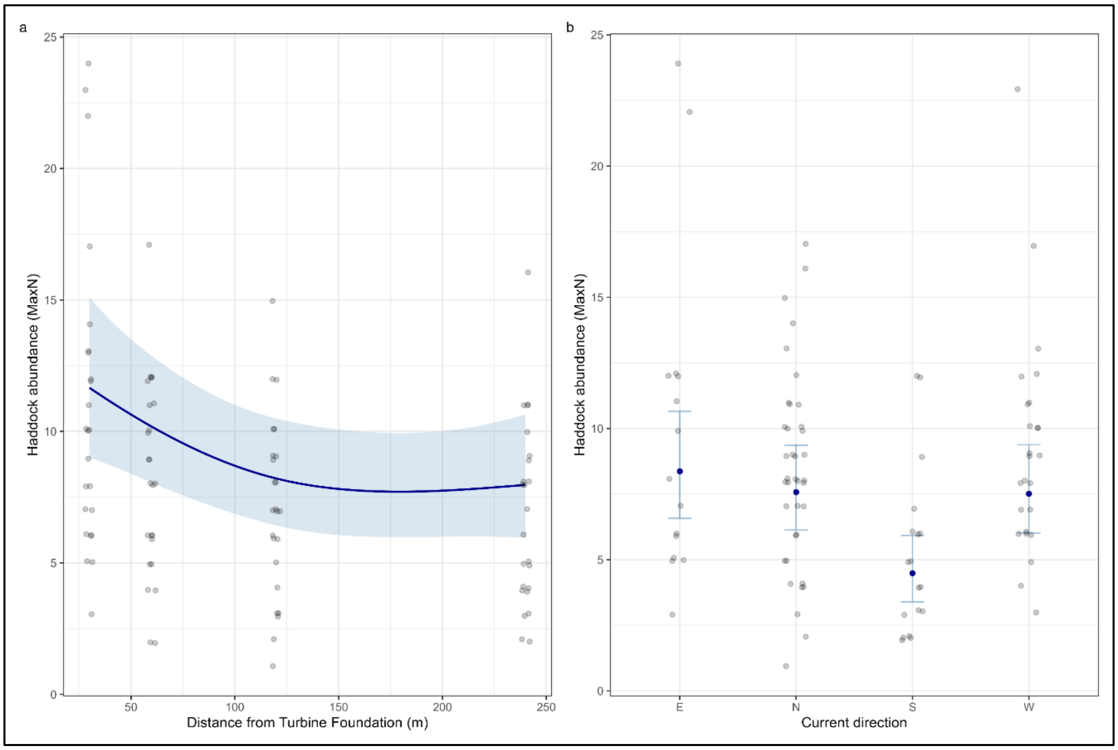


**Supplementary Figure 9.** Plots of raw data and estimated values with standard errors (light blue shaded ribbon or error bars) for significant terms from the haddock abundance GAM.

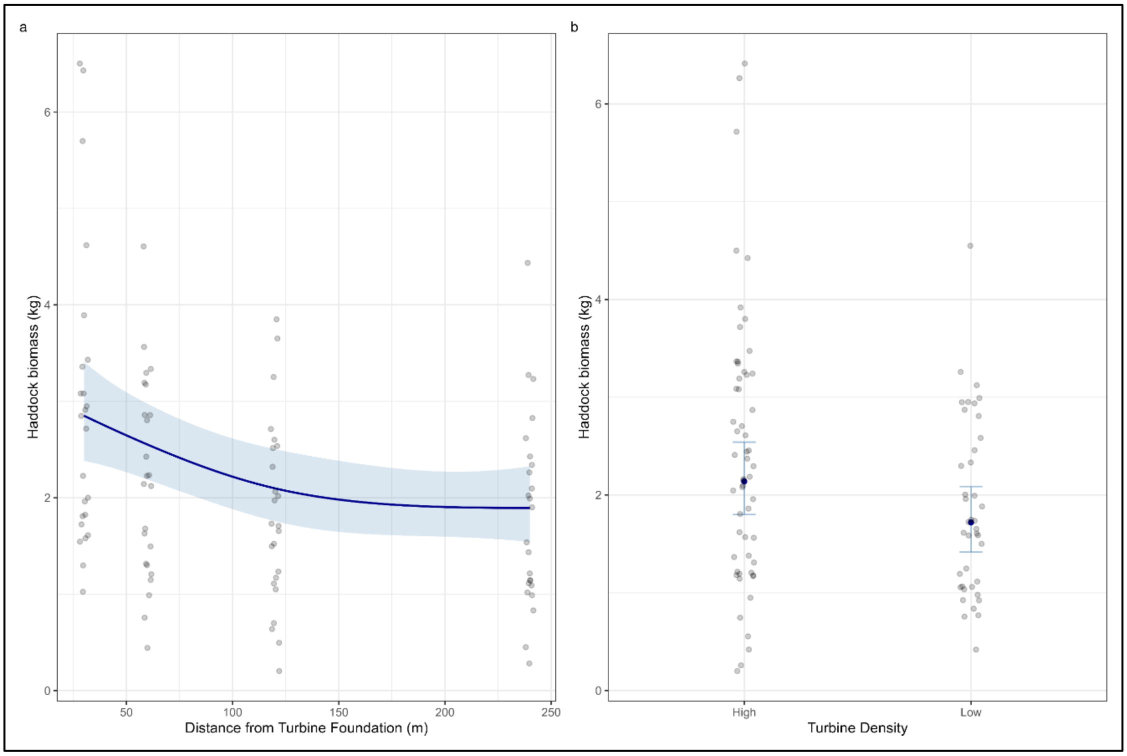


**Supplementary Figure 20.** Plots of raw data and estimated values with standard errors (light blue shaded ribbon or error bars) for significant terms from the haddock biomass GAM.

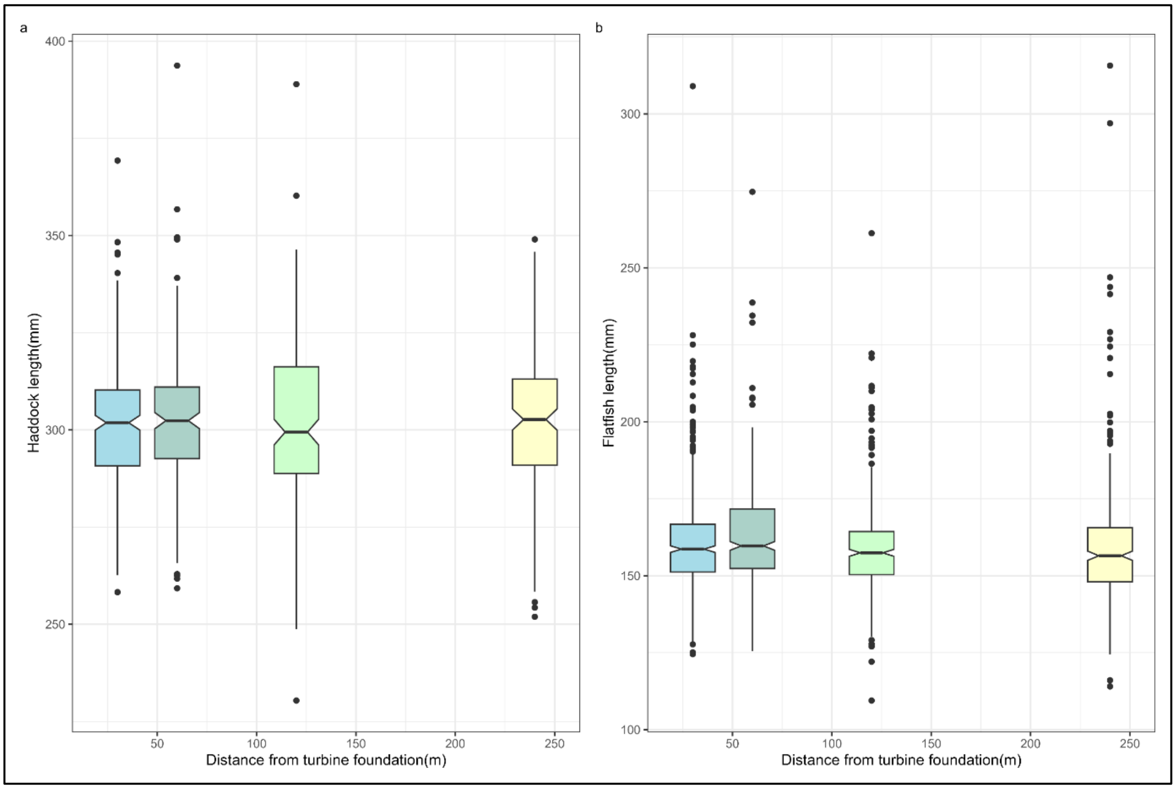


**Supplementary Figure 31.** Box and whisker plots of raw haddock (**a**) and flatfish (**b**) length data from Beatrice BRUV surveys.

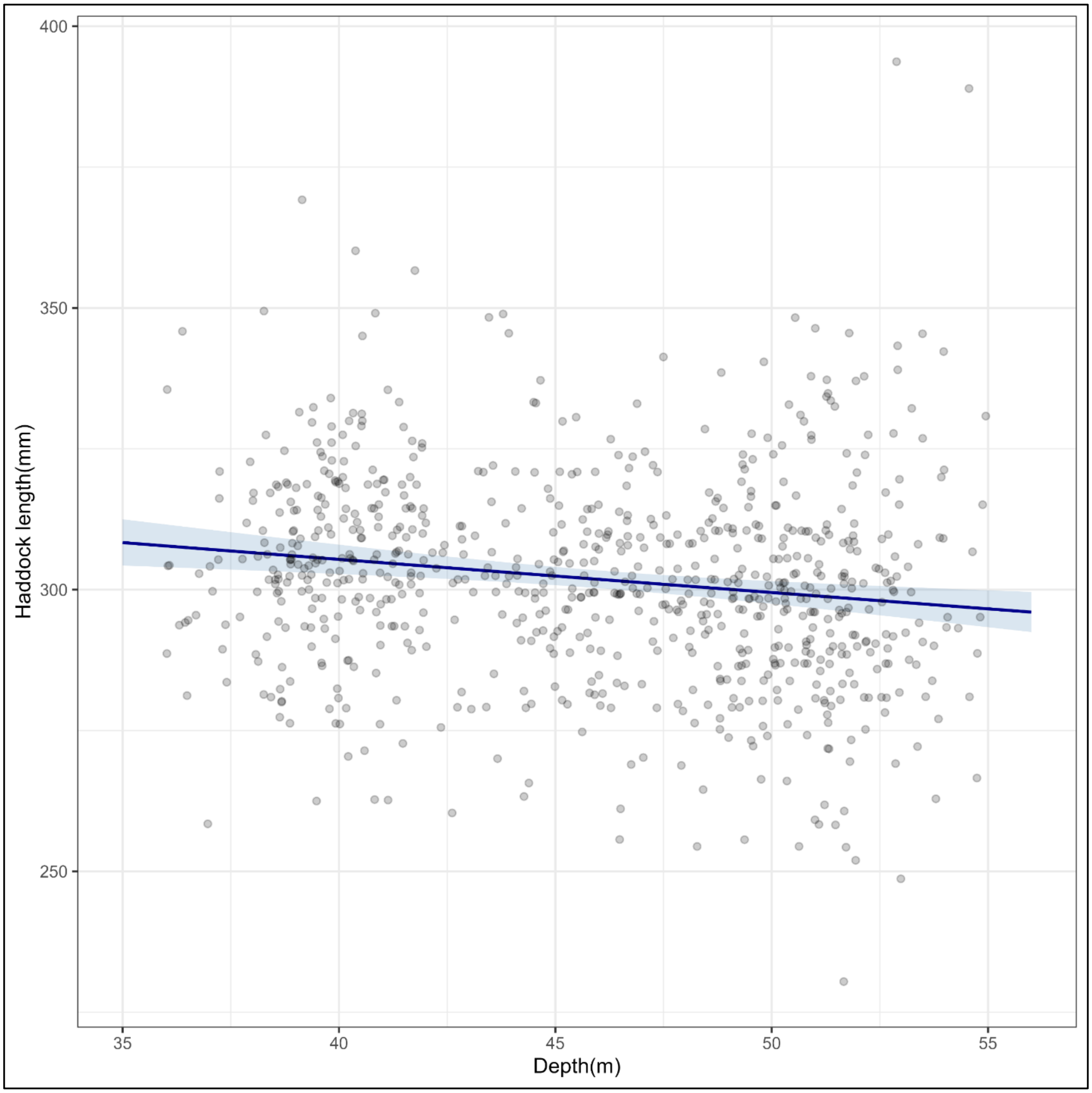


**Supplementary Figure 42.** Plot of raw data and estimated values with standard errors of significant depth term from the haddock length GLMM model.

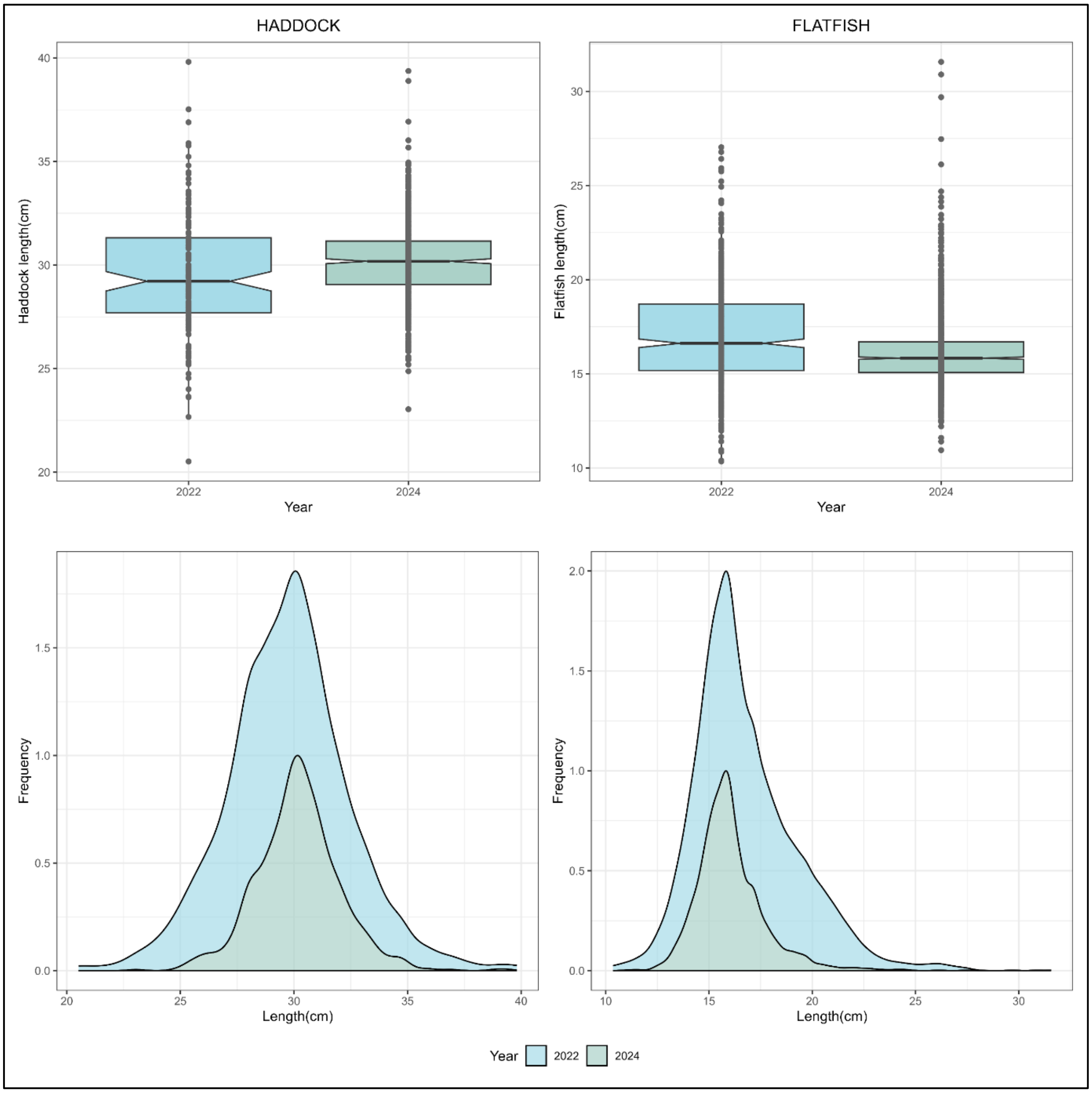


**Supplementary Figure 53.** Plot of raw data box plots and frequency histograms of haddock and flatfish length for fish sampled at 30m from the turbine foundations in Beatrice wind farm in 2022 and 2024.

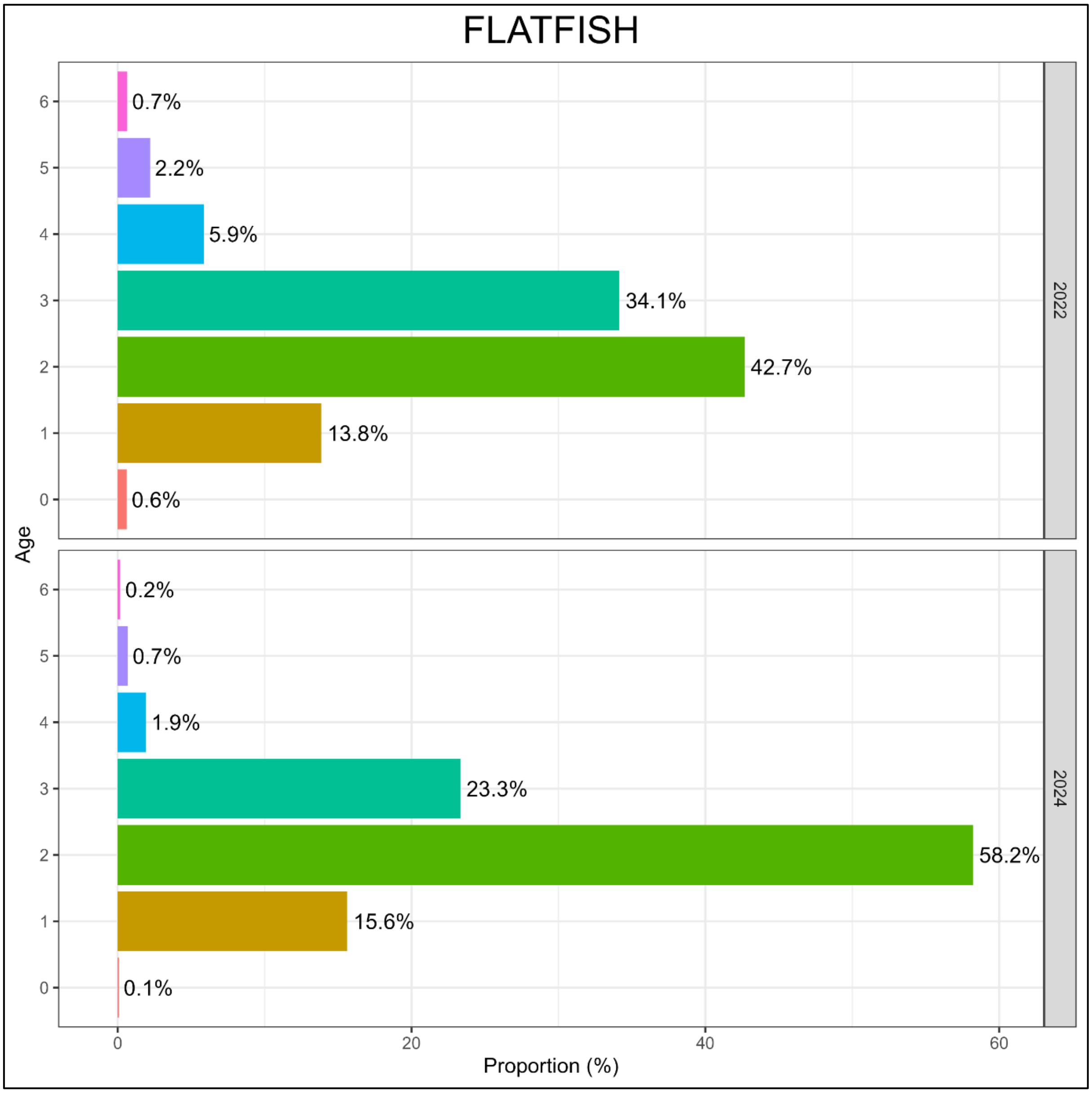


**Supplementary Figure 64.** Plot of estimated age classes and proportions for flatfish sampled at ~30m from the turbine foundations in Beatrice wind farm in 2022 and 2024.

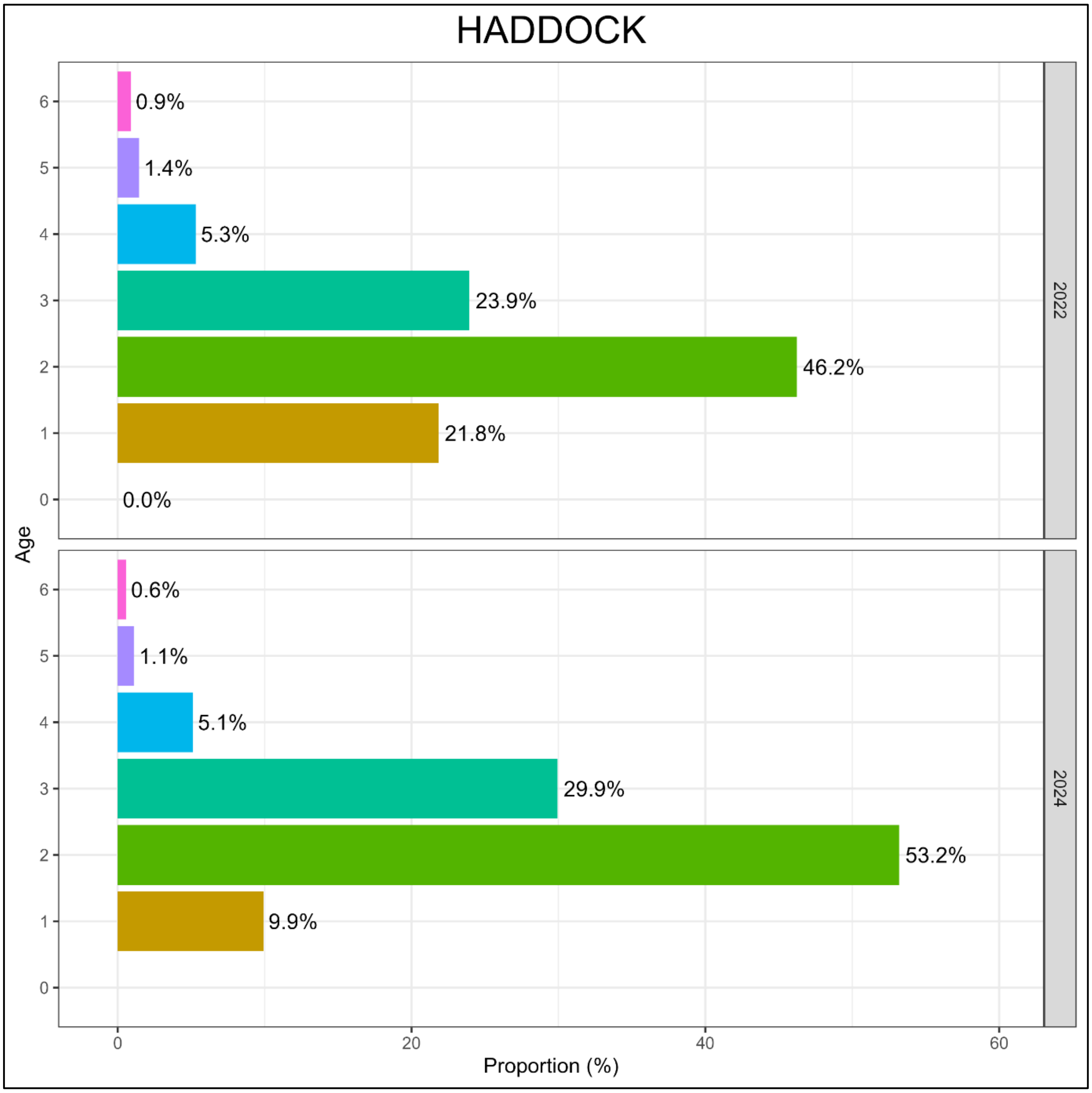


**Supplementary Figure 75.** Plot of estimated age classes and proportions for haddock sampled at ~30m from the turbine foundations in Beatrice wind farm in 2022 and 2024

### 1.2 Supplementary Tables

**Supplementary Table 8.** YOLO detection model performance statistics on a validation set of 199 images sampled from BRUV survey data collected at the study site in 2022. A total of 166 flatfish and 152 haddock were present in the ground truth labels. The metrics reported are: precision = the proportion of predicted instances that were present in the ground truth data; recall = the proportion of ground truth instances successfully detected; mAP = the area under the precision-recall curve, showing the average model precision over varying confidence thresholds and across object classes.

| **Class** | **Precision** | **Recall** | **mAP** |
|---|---|---|---|
| Flatfish (*Pleuronectiformes spp.*) | 0.801 | 0.819 | 0.849 |
| Haddock (*Melanogrammus aeglefinus*) | 0.878 | 0.866 | 0.903 |
| All | 0.839 | 0.842 | 0.876 |

**Supplementary Table 2.** Best fitting GLMM for haddock length. Terms with significant p-values below the 0.05 level are shown in bold.

| | **Haddock length** | | |
|---|---|---|---|
| ***Fixed terms*** | ***Estimates*** | ***SE*** | ***p-values*** |
| (Intercept) | 5.799 | 0.026 | **<0.001** |
| Depth | -0.002 | 0.001 | **0.001** |
| **Random Effects** | | | |
| $\sigma^2$ | 0.0035 | | |
| $\tau_{00\ WTG}$ | 0.0001 | | |
| ICC | 0.0150 | | |
| $N_{WTG}$ | 24 | | |
| Observations | 788 | | |
| Marginal $R^2$ / Conditional $R^2$ | 0.024 / 0.038 | | |

### 1.3 Supplementary Videos

**Supplementary Video 1** Example detections from the YOLO model used for Haddock and Flatfish MaxN estimation. The model was used to detect haddock and flatfish in real time with a confidence threshold of 0.3. All detections are shown with their class label and confidence score.
